## Supplementary Materials for "Clinically applicable histopathological diagnosis system for gastric cancer detection using deep learning"

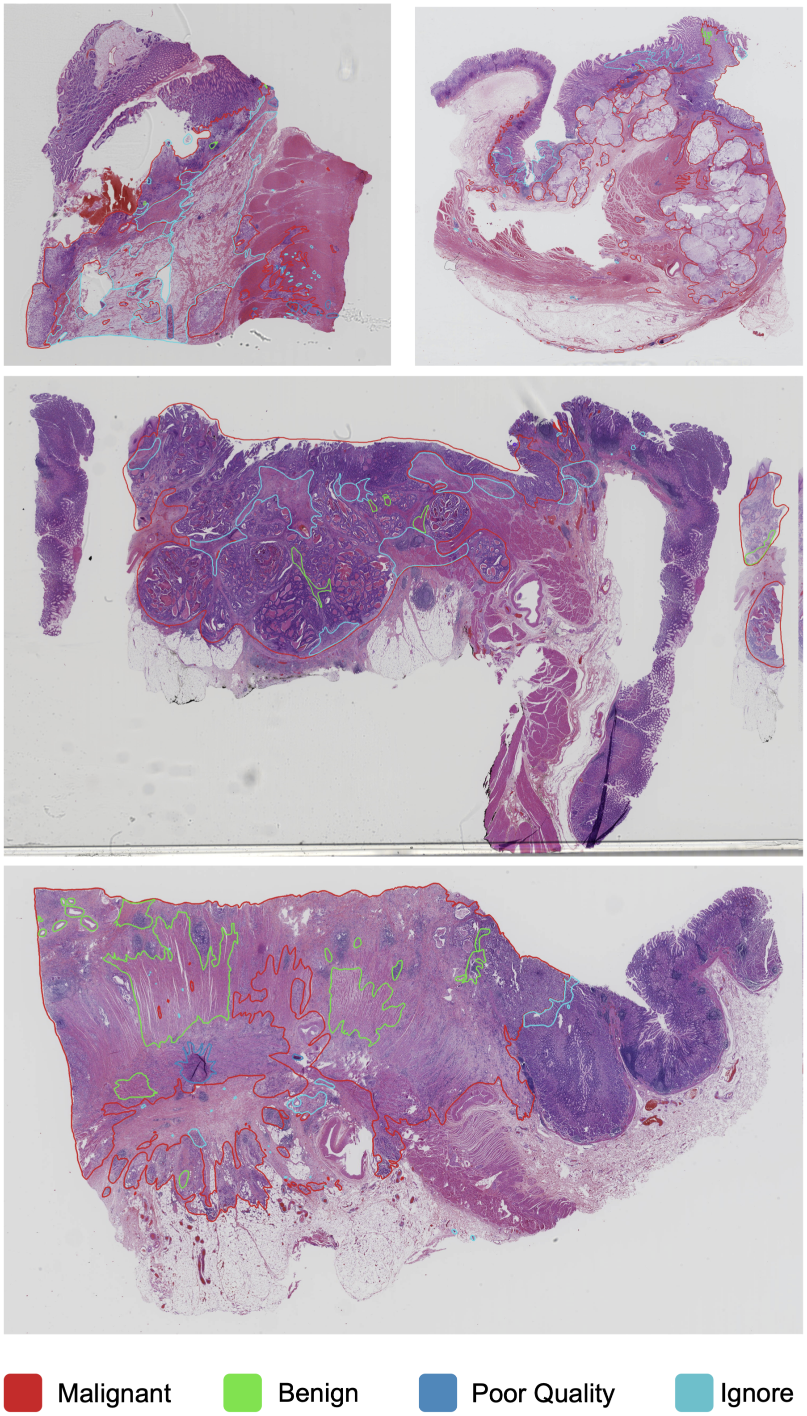


Figure S1: **Four examples of labelled WSIs.**

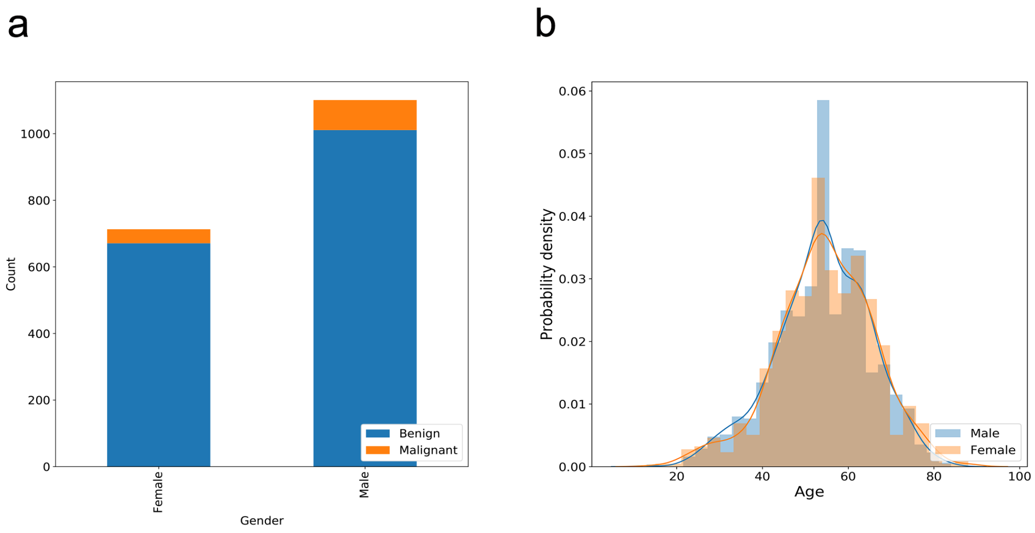


Figure S2: **Patient-level data distribution of the daily gastric dataset.** **a**, Patient gender distribution. **b**, Patient age distribution.

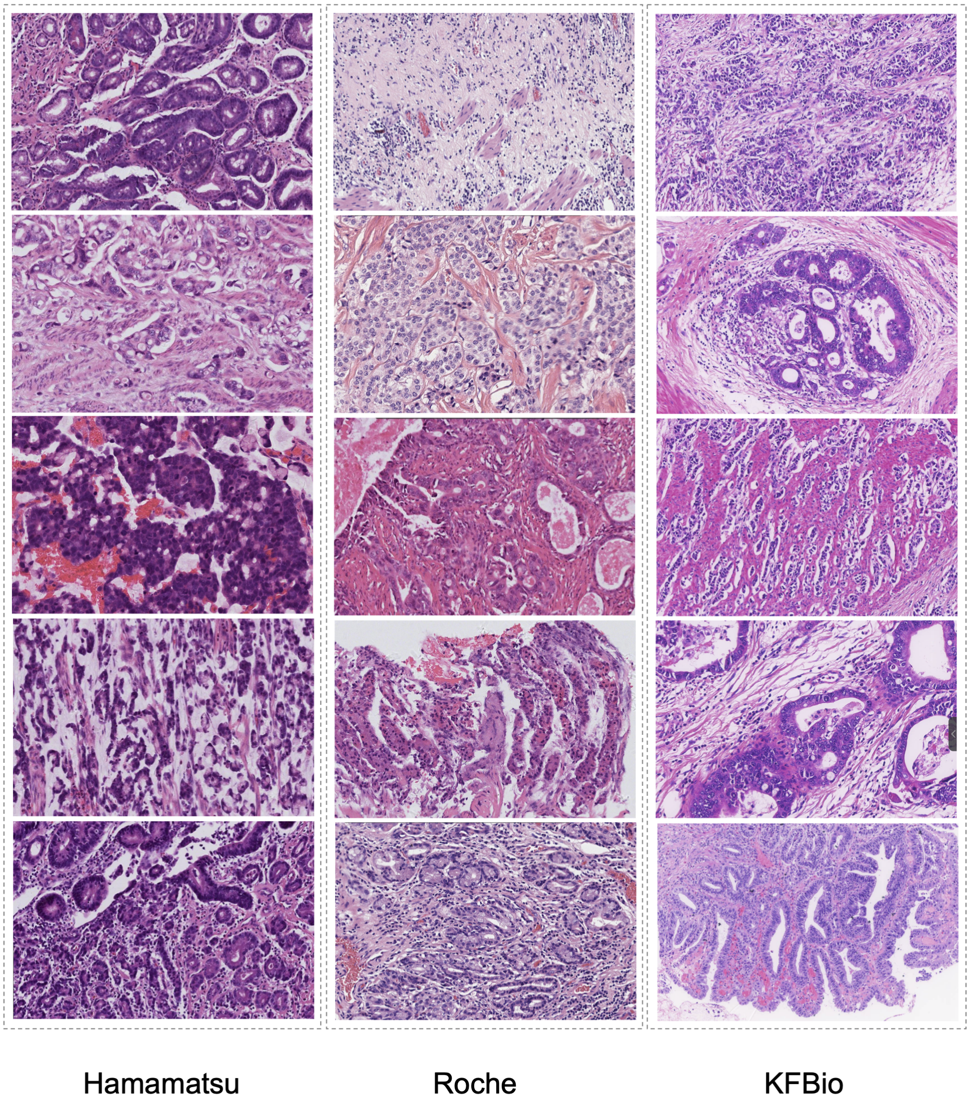


Figure S3: **Example WSIs digitalized by three different scanners from the daily gastric dataset.** All the images were captured with a 20× objective.

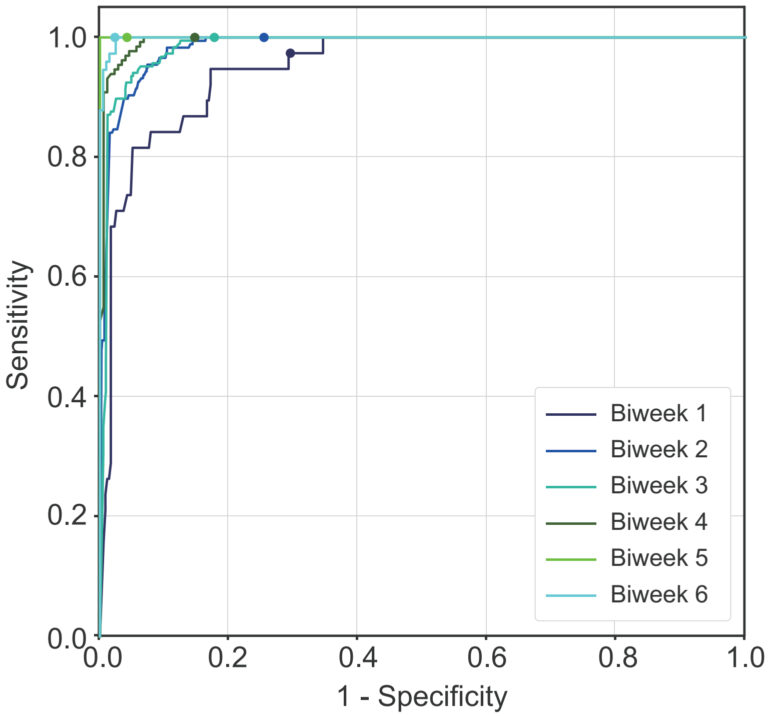


Figure S4: **Model performance on the daily gastric dataset.**

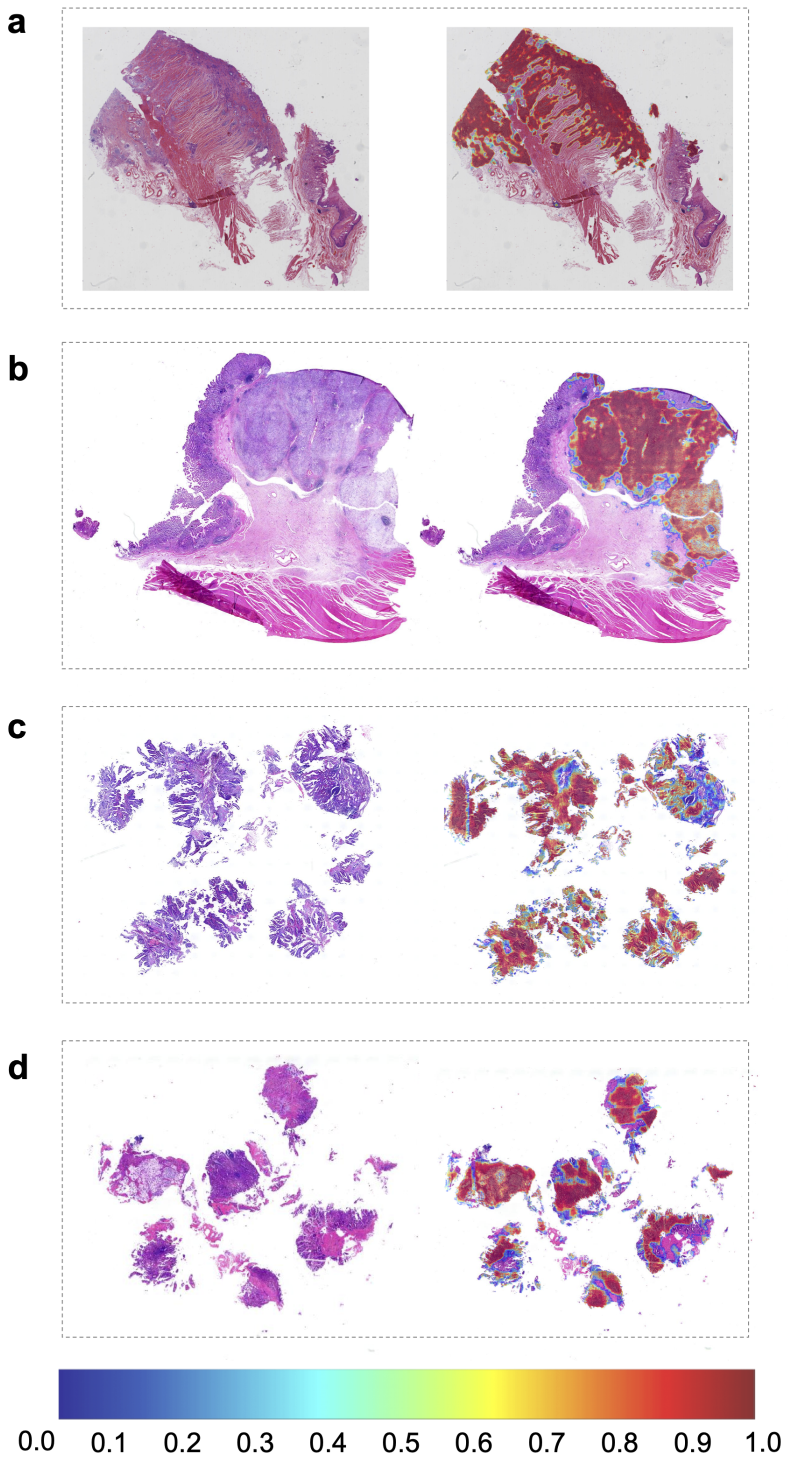


Figure S5: **Four examples of deep learning model predictions in the form of heatmaps.**

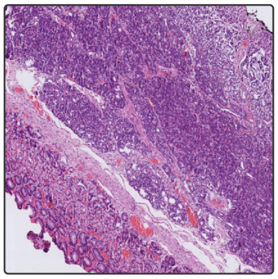

(a) Heterotopic pancreas

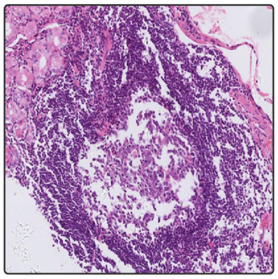

(b) Distorted lymphoid follicle

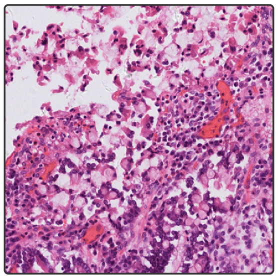

(c) Disorientated and sloughing foveolar epithelium

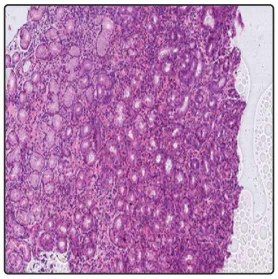

(d) Trapped air bubbles

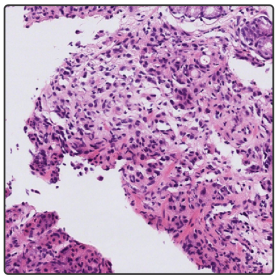

(e) Inflamed granulation tissue

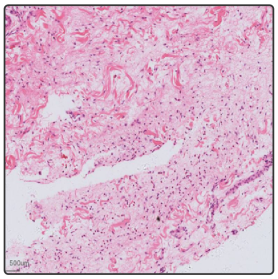

(f) Fibrinoinflammatory exudates

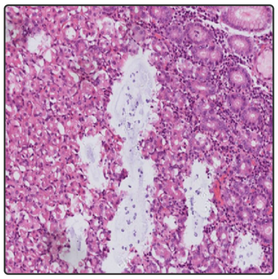

(g) Shreddered section

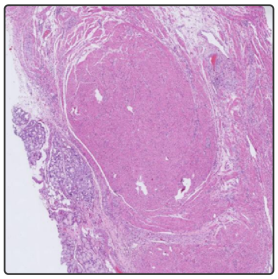

(h) Carcinoma with co-existing stromal tumor

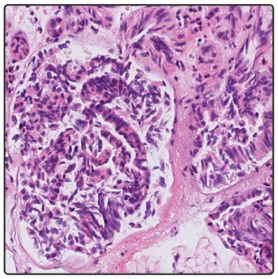

(i) Crushed tissue

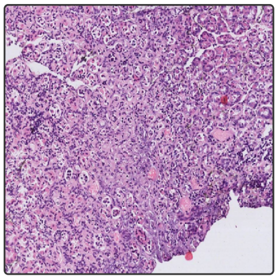

(j) Thick section

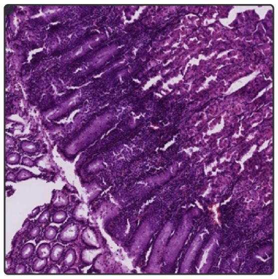

(k) Overstained section

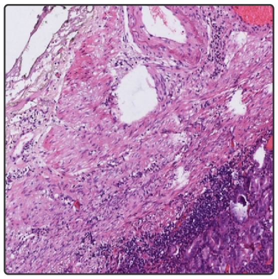

(l) Inflammatory infiltrates in muscularis propria

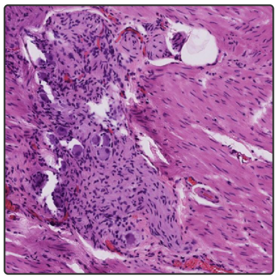

(m) Hypertrophic myenteric plexus

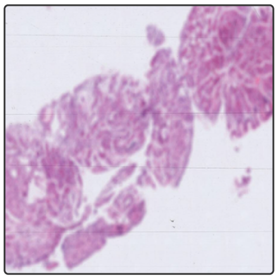

(n) Defocused section

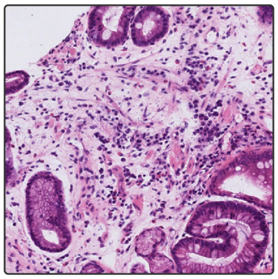

(o) Inflammatory infiltrates in edematous stroma

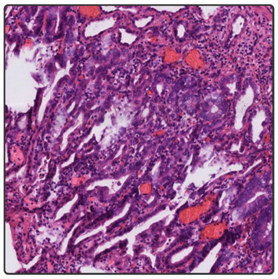

(p) Regenerative atypia accompanied by marked inflammation

Figure S6: **More false-positive cases.**

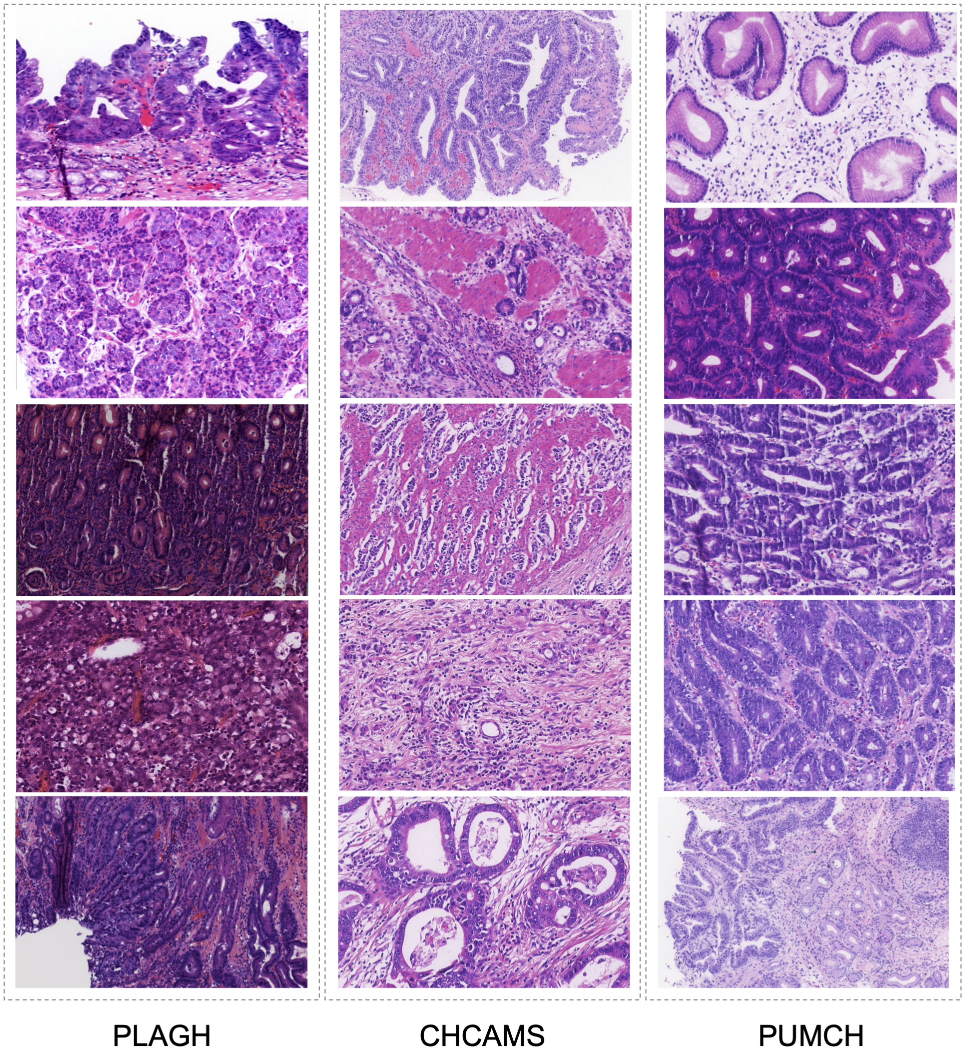


Figure S7: **Example WSIs with visual difference digitalized by KF-Pro-005 from three hospitals.** PLAGH and PUMCH used automatic H&E staining with Leica AutoStainer XL in practice, while CHCAMS adopted automatic H&E staining with Roche Ventana HE 600. Significant visual difference could be found in the images. All the images were captured with a 20× objective.

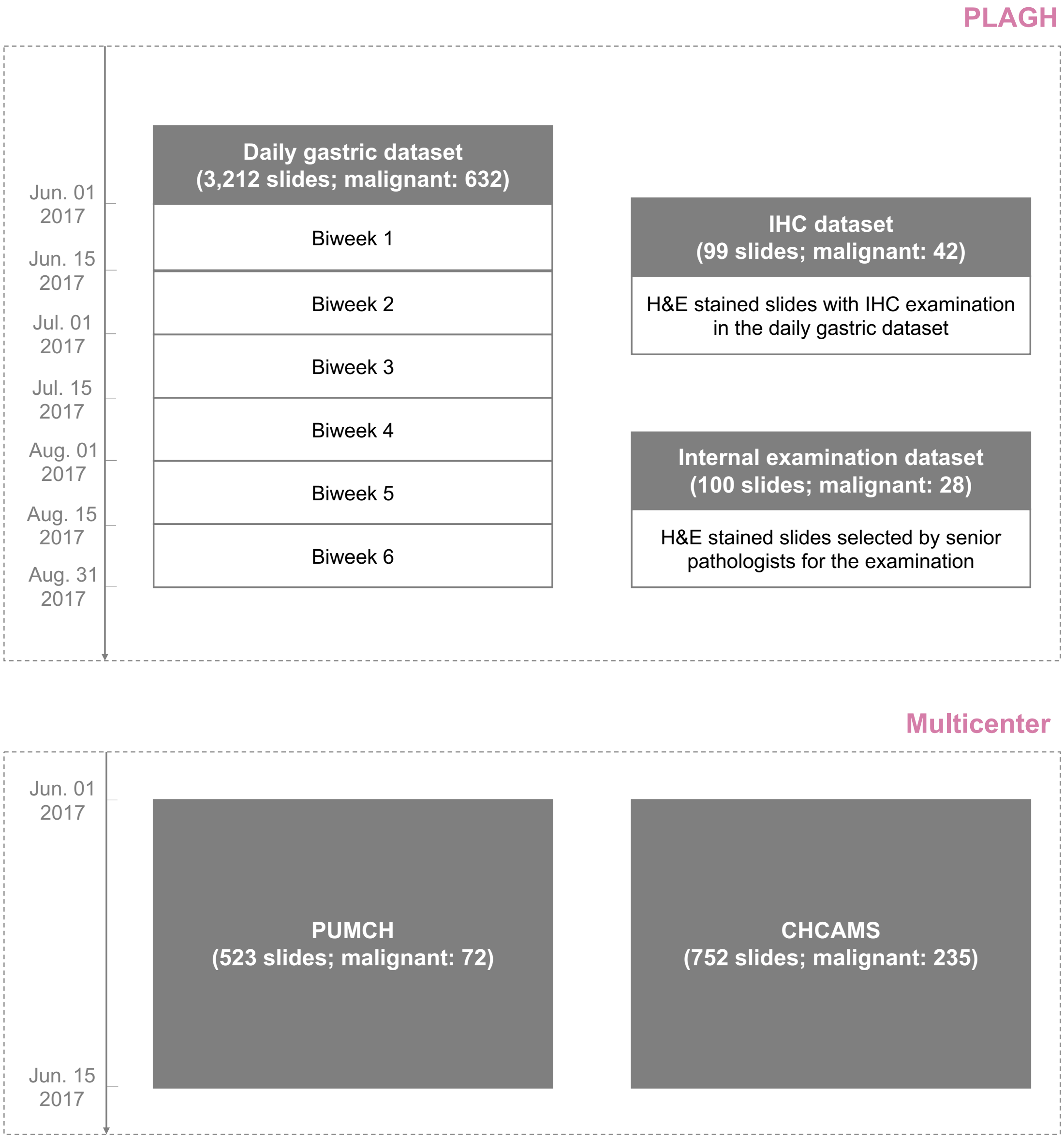


Figure S8: **Illustration of the test datasets.**

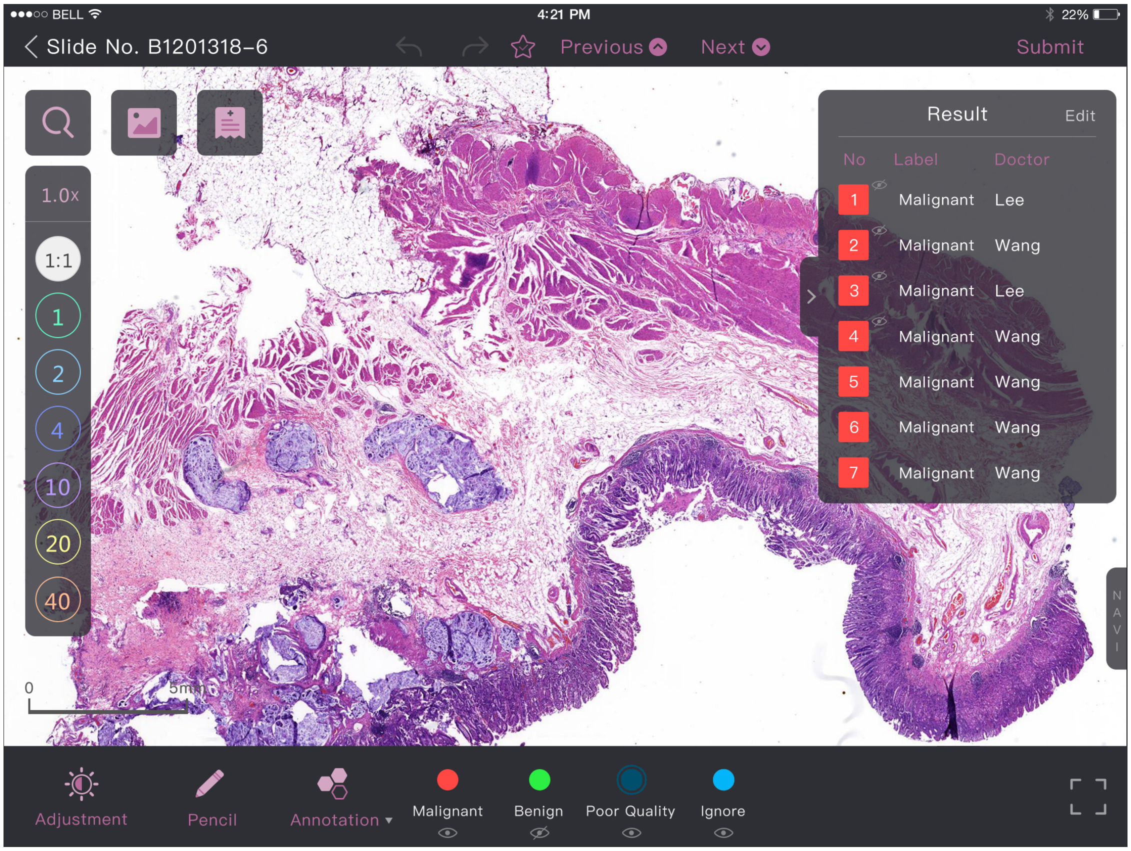


Figure S9: **iPad-based annotation system interface.**

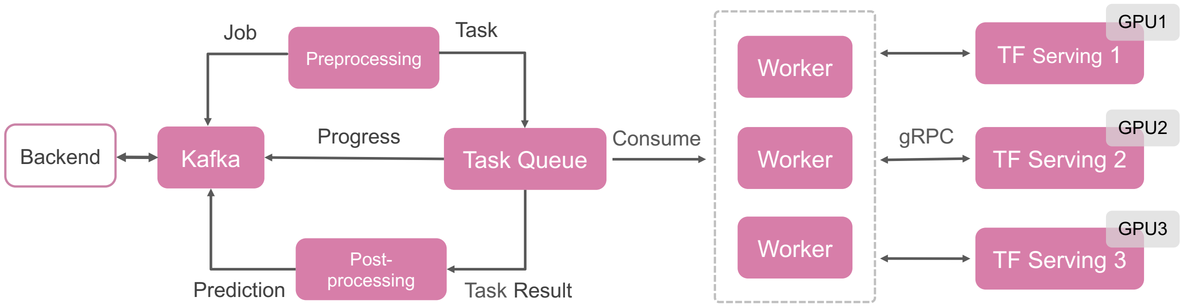


Figure S10: **AI assistance system architecture.**

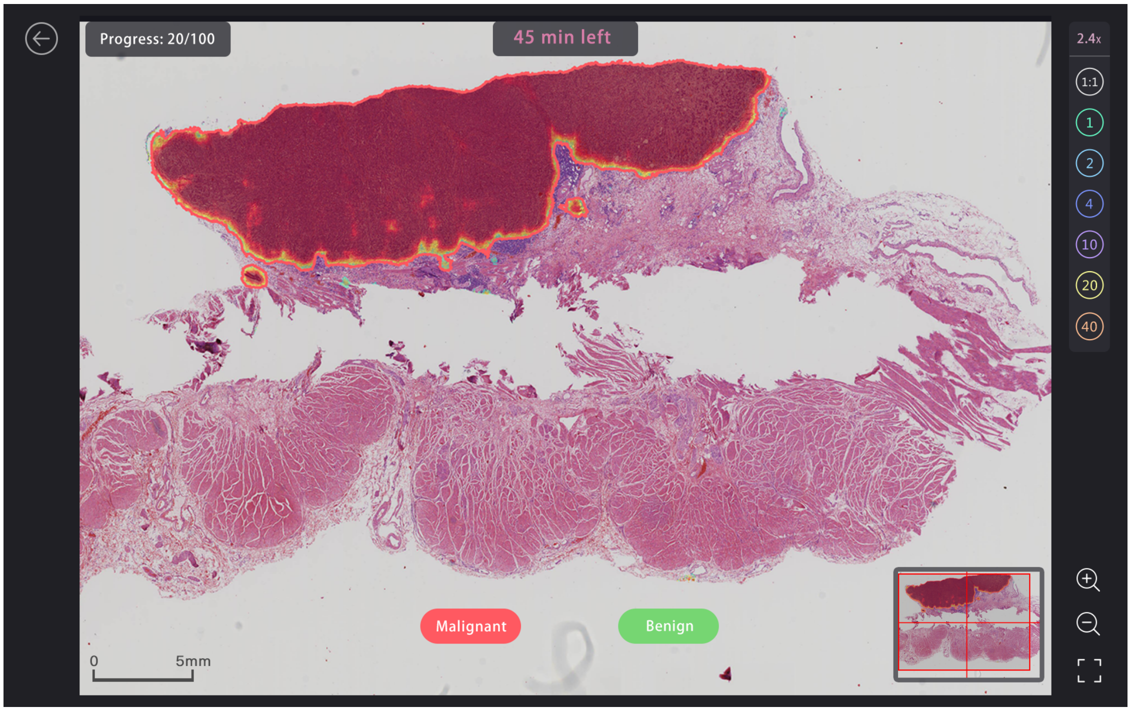


Figure S11: **The interface of the AI assistance system used in the internal examination.**

Table S1: **Details of annotation pathologists.**

| Name | Position | Education | Experience (year) | Role |
|---|---|---|---|---|
| Huaiyin Shi | Chief Pathologist | PhD | 31 | Final check |
| Zhigang Song | Associate Chief Pathologist | Master | 16 | Final check |
| Zhanbo Wang | Associate Chief Pathologist | Master | 16 | Labelling |
| Jing Yuan | Associate Chief Pathologist | PhD | 16 | Labelling |
| Chunkai Yu | Associate Chief Pathologist | PhD | 16 | Labelling |
| Yong Huang | Attending Pathologist | PhD | 19 | Labelling |
| Jinhong Liu | Attending Pathologist | PhD | 16 | Labelling |
| Xiaohui Ding | Attending Pathologist | Master | 16 | Labelling |
| Xin Chen | Attending Pathologist | Master | 7 | Labelling |
| Wei Jin | Attending Pathologist | Master | 7 | Labelling |
| Xiangnan Gou | Attending Pathologist | Master | 7 | Labelling |
| Liwei Shao | Attending Pathologist | Master | 4 | Labelling |

Table S2: **Abbreviations of tumour subtypes.**

| | |
|---|---|
| HGIN | High grade intraepithelial neoplasia |
| TAC | Tubular adenocarcinoma |
| MucAC | Mucinous adenocarcinoma |
| PCC | Poorly cohesive carcinoma |
| MixAC | Mixed adenocarcinoma |

Table S3: **Performance of different slide-level prediction approaches on the validation dataset.**

| Predictor | AUC | Sensitivity | Specificity | Accuracy |
|---|---|---|---|---|
| Random forest | 0.970 | 0.986 | 0.520 | 0.723 |
| Top 10 probabilities | 0.983 | 0.986 | 0.532 | 0.753 |
| Top 100 probabilities | 0.983 | 0.986 | 0.571 | 0.773 |
| Top 200 probabilities | 0.982 | 0.986 | 0.604 | 0.790 |
| Top 500 probabilities | 0.985 | 0.986 | 0.636 | 0.807 |
| Top 1,000 probabilities | 0.988 | 0.986 | 0.740 | 0.860 |
| Top 2,000 probabilities | 0.983 | 0.986 | 0.734 | 0.857 |
| Top 5,000 probabilities | 0.979 | 0.986 | 0.721 | 0.850 |
| Top 10,000 probabilities | 0.976 | 0.986 | 0.675 | 0.827 |

Table S4: **Performance of different classification and segmentation models for patch-level classification on the validation dataset.**

| Deep learning model | AUC | Sensitivity | Specificity | Accuracy |
|---|---|---|---|---|
| ResNet-50 | 0.853 | 0.779 | 0.777 | 0.778 |
| Inception v3 | 0.887 | 0.864 | 0.765 | 0.806 |
| DenseNet | 0.834 | 0.804 | 0.713 | 0.750 |
| U-Net | 0.779 | 0.950 | 0.369 | 0.550 |
| DeepLab v2 | 0.884 | 0.918 | 0.710 | 0.775 |
| DeepLab v3 | 0.945 | 0.944 | 0.780 | 0.833 |

Table S5: **Model performance on the daily gastric WSI digitalized by three scanners.**

| Digital scanner | Total | Malignant | Benign | AUC | Accuracy | Sensitivity | Specificity |
|---|---|---|---|---|---|---|---|
| KF-PRO-005 | 403 | 80 | 323 | 0.995 | 0.950 | 1.0 | 0.938 |
| Hamamatsu NanoZoomer S360 | 1832 | 352 | 1480 | 0.983 | 0.781 | 1.0 | 0.728 |
| Ventana DP200 | 977 | 200 | 777 | 0.992 | 0.918 | 0.995 | 0.898 |

Table S6: **Pathologists' performance in the trainees' examination with one hour constraint.**

| Group name | Pathologist ID | Individual performance | | | | Group performance (average) | | | |
|---|---|---|---|---|---|---|---|---|---|
| | | Acc. | Sen. | Spec. | Time (min) | Acc. | Sen. | Spec. | Time (min) |
| Microscope | 6 | 0.820 | 0.929 | 0.778 | 44 | 0.850 | 0.821 | 0.861 | 53.30 |
| | 8 | 0.870 | 0.786 | 0.903 | 51 | | | | |
| | 9 | 0.830 | 0.643 | 0.903 | 72 | | | | |
| | 10 | 0.880 | 0.929 | 0.861 | 47 | | | | |
| Digital | 0 | 0.820 | 0.429 | 0.972 | 48 | 0.845 | 0.696 | 0.903 | 48.25 |
| | 2 | 0.860 | 0.750 | 0.903 | 45 | | | | |
| | 11 | 0.860 | 0.857 | 0.861 | 52 | | | | |
| | 1 | 0.840 | 0.750 | 0.875 | 48 | | | | |
| AI | 5 | 0.910 | 0.786 | 0.958 | 41 | 0.858 | 0.839 | 0.865 | 45.25 |
| | 4 | 0.850 | 0.786 | 0.875 | 43 | | | | |
| | 3 | 0.840 | 0.964 | 0.792 | 47 | | | | |
| | 7 | 0.830 | 0.821 | 0.833 | 50 | | | | |

Table S7: **Pathologists' performance in the trainees' examination without time constraint.**

| Group name | Pathologist ID | Individual performance | | | | Group performance (average) | | | |
|---|---|---|---|---|---|---|---|---|---|
| | | Acc. | Sen. | Spec. | Time (min) | Acc. | Sen. | Spec. | Time (min) |
| Microscope | 1 | 0.820 | 0.929 | 0.778 | 60 | 0.810 | 0.911 | 0.771 | 59.25 |
| | 2 | 0.880 | 0.893 | 0.875 | 75 | | | | |
| | 7 | 0.810 | 0.964 | 0.750 | 60 | | | | |
| | 11 | 0.730 | 0.857 | 0.681 | 42 | | | | |
| Digital | 8 | 0.820 | 0.821 | 0.819 | 30 | 0.853 | 0.857 | 0.851 | 40 |
| | 5 | 0.910 | 0.821 | 0.944 | 46 | | | | |
| | 4 | 0.840 | 0.893 | 0.819 | 41 | | | | |
| | 3 | 0.840 | 0.893 | 0.819 | 43 | | | | |
| AI | 6 | 0.860 | 0.893 | 0.847 | 61 | 0.870 | 0.813 | 0.892 | 50.75 |
| | 9 | 0.840 | 0.679 | 0.903 | 31 | | | | |
| | 10 | 0.920 | 0.857 | 0.944 | 53 | | | | |
| | 0 | 0.860 | 0.821 | 0.875 | 58 | | | | |

Table S8: **Distribution of benign and malignant cases and tumour subtypes in the datasets by individual patient.**

| Dataset | Specimen | Benign | Malignant | Tumour subtype | | | | |
|---|---|---|---|---|---|---|---|---|
| | | | | HGIN | TAC | MucAC | PCC | MixAC |
| Training | Biopsy | 440 | 102 | 22 | 59 | 21 | 0 | 0 |
| | Surgical | 50 | 908 | 151 | 579 | 176 | 6 | 51 |
| Validation | Biopsy | 119 | 60 | 0 | 27 | 6 | 13 | 14 |
| | Surgical | 7 | 86 | 7 | 51 | 8 | 12 | 15 |
| Internal examination | Biopsy | 68 | 27 | 3 | 20 | 0 | 7 | 0 |
| | Surgical | 4 | 1 | 0 | 1 | 0 | 0 | 0 |
| IHC | Biopsy | 30 | 12 | 0 | 11 | 0 | 1 | 0 |
| | Surgical | 1 | 9 | 0 | 9 | 0 | 0 | 0 |
| Daily gastric (PLAGH) | Biopsy | 1599 | 61 | 12 | 42 | 4 | 2 | 10 |
| | Surgical | 36 | 118 | 16 | 83 | 6 | 4 | 25 |
| Multicentre (PUMCH) | Biopsy | 330 | 14 | 5 | 10 | 0 | 0 | 0 |
| | Surgical | 3 | 8 | 0 | 6 | 0 | 0 | 2 |
| Multicentre (CHCAMS) | Biopsy | 396 | 59 | 6 | 51 | 0 | 3 | 5 |
| | Surgical | 15 | 71 | 2 | 49 | 1 | 3 | 19 |

Table S9: **Distribution of benign and malignant cases and tumour subtypes in the datasets by individual slide.**

| Dataset | Specimen | Benign | Malignant | Tumour subtype | | | | |
|---|---|---|---|---|---|---|---|---|
| | | | | HGIN | TAC | MucAC | PCC | MixAC |
| Training | Biopsy | 639 | 102 | 22 | 59 | 21 | 0 | 0 |
| | Surgical | 93 | 1,289 | 242 | 760 | 274 | 11 | 63 |
| Validation | Biopsy | 143 | 60 | 0 | 27 | 6 | 13 | 14 |
| | Surgical | 11 | 86 | 5 | 51 | 8 | 12 | 15 |
| Internal examination | Biopsy | 68 | 27 | 3 | 20 | 0 | 7 | 0 |
| | Surgical | 4 | 1 | 0 | 1 | 0 | 0 | 0 |
| IHC | Biopsy | 49 | 14 | 0 | 13 | 0 | 1 | 0 |
| | Surgical | 8 | 28 | 0 | 28 | 0 | 0 | 0 |
| Daily gastric (PLAGH) | Biopsy | 2,087 | 124 | 20 | 93 | 5 | 8 | 6 |
| | Surgical | 495 | 508 | 51 | 354 | 26 | 22 | 118 |
| Multicentre (PUMCH) | Biopsy | 496 | 30 | 7 | 24 | 0 | 0 | 0 |
| | Surgical | 27 | 42 | 0 | 38 | 0 | 0 | 4 |
| Multicentre (CHCAMS) | Biopsy | 685 | 81 | 20 | 68 | 0 | 5 | 8 |
| | Surgical | 67 | 154 | 6 | 105 | 3 | 6 | 43 |

Table S10: **Features used for the random forest model.** The eccentricity, extend, major axis length, and solidity are defined as ellipse with the same second moment, ratio of the region area over the bounding box, length of the major axis of the ellipse with the same normalized second central moment, and ratio of the region area over the surrounding convex, respectively.

| Number | Feature definition |
|---|---|
| 1-5 | Ratios of cancer to tissue (thresholds: 0.5, 0.6, 0.7, 0.8, 0.9) |
| 6-10 | Ratios of probability sum of cancer to tissue (thresholds: 0.5, 0.6, 0.7, 0.8, 0.9) |
| 11-14 | Largest area, eccentricity, extend, and bounding box area |
| 15 | Major axis length |
| 16-17 | Maximum/minimum probability in the region |
| 18 | Largest mean probability in the region |
| 19 | Aspect ratio of the bounding box |
| 20 | Solidity |
| 21-24 | Second largest area, eccentricity, extend, and bounding box area |
| 25 | Minor axis length |
| 26-27 | Second maximum/minimum probability in the region |
| 28 | Second largest mean probability in the region |
| 29 | Aspect ratio of the bounding box (second largest) |
| 30 | Second largest solidity |
